# Development of a Stabilized Alginate-based Hydrogel for Oral Delivery of Encapsulated Live Cultures and Allowing their Intact Passage Through the Digestive System

**DOI:** 10.64898/2026.01.31.703036

**Authors:** Bruce J. Godfrey, Pei-Hsin Wang, Prakit Saingam, Happy Tju, Mari-Karoliina Henriika Winkler

## Abstract

Alginate hydrogels are widely used for biocompatible encapsulation due to their low cost, mild gelation conditions, and scalability; however, their limited mechanical strength and poor chemical stability under physiological conditions restrict their utility for oral delivery applications. In particular, the development of robust alginate formulations capable of surviving gastrointestinal salt and pH exposures is critical for advancing encapsulated microbial therapeutics for chronic kidney disease (CKD). In this study, we investigated the incorporation of ferric iron into calcium alginate networks as a strategy to enhance gel stability while maintaining biocompatibility. Using a three-ion competition approach, we achieved controlled introduction of ferric ions into calcium alginate gels without significantly altering bulk mechanical properties relative to standard calcium alginate. Although the initial ferric-containing gels displayed comparable modulus and structure, post-treatment with chitosan under mildly acidic conditions produced a dramatic increase in gel stability in physiological salt concentrations across both acidic and neutral pH environments. Ferric-containing gels formed at pH 4.6 absorbed negligible chitosan, in contrast to iron-free alginate gels, which incorporated substantial chitosan under identical conditions. These results support the formation of a thin, dense interfacial complex between chitosan, ferric ions, and alginate at the gel surface, which reinforces the matrix and inhibits dissolution. The resulting hybrid ferric–calcium alginate formulation enabled the production of sub-millimeter beads capable of encapsulating live *Thauera aminoaromatica* while preserving anaerobic p-cresol degradation activity at 37 °C using nitrate as an electron acceptor. Collectively, these findings establish ferric-modified alginate hydrogels as a promising, scalable platform for stable oral delivery of encapsulated microbial therapeutics.

## Introduction

Uremic toxins such as cresyl sulfate and indoxyl sulfate are generated in the liver from bacteria precursors p-cresol and indole, which are produced by enteric bacteria in the colon through amino acids metabolism. These precursors are absorbed into the bloodstream and subsequently converted by hepatic sulfonation [1]. The resulting protein-bound uremic toxins (PBUT) are tightly bind to serum albumin and are ultimately clear via renal excretion. However, these protein-bound uremic toxins are not efficiently removed by damaged kidneys or by dialysis so these toxins accumulate systemically in patients with chronic kidney disease (CKD) [1, 2], resulting in persistently elevated plasma concentrations that correlate with disease severity and adverse outcomes [3]. There is no treatment currently available which can effectively prevent the accumulation of PBUTs. It would be desirable to harness the biochemical specificity and metabolic diversity of bacteria to reduce the production of toxic metabolites in the human gut by the existing microbiome without directly altering the microbiome and risking unintended consequences of such alteration. Such an approach could provide the basis of a novel treatment useful for patients across multiple stages in CKD progression.

To enable this treatment modality, live microbial therapeutics capable of metabolizing uremic toxins or their precursors could be encapsulated in hydrogel particles. Encapsulation would allow the microbes to remain metabolically active while being physically retained within the gel matrix, thereby minimizing direct interaction with and unintended alterations of the native gut microbiome [4]. Archiving this requires a hydrogel system that can be solidified under mild conditions that do not reduce the viability of the encapsulated cells and which is stable enough to survive exposure to gastric acid, digestive enzymes, extended contact with physiological salt concentrations at pHs ranging from ∼5 to ∼8, and a variety of actively growing microbes that may be encountered at various points in the intestinal tract of mammals [5]. The hydrogel must be composed of materials approved for human consumption and it must be practical to produce in the form of sub-millimeter particles. This is important both due to the requirements of feeding rodents by gavage during development testing and to minimize fluid volumes required during use by patients. For a therapeutic intended for repeated oral administration, minimizing the total volumetric intake is important to maintain patient compliance and limit fluid burden. Smaller hydrogel particles, with their increased surface-to-volume ratio, can enhance substrate exchange and microbial activity, thereby maximizing toxin precursor turnover per administered volume [6]. This consideration is particularly relevant in chronic kidney disease, where fluid intake is often restricted to prevent fluid overload and associated complications due to impaired renal excretion [7].

A range of natural and synthetic gel-forming polymers can satisfy several of the key requirements for hydrogel fabrication including alginate, polyvinyl alcohol (PVA), and polyethylene glycol (PEG) acrylates [8, 9]. PVA hydrogels exhibit strong mechanical performance and good biocompatibility but the high viscosity of the starting solutions complicates the generation of uniform and small particle[10]. PEG acrylate and similar vinylic polymer systems offer tunable network properties but typically require free radical initiators that tend to have some degree of toxicity towards live cells and the water soluble monomers are themselves toxic and impose the requirement of using an immiscible oil phase to allow droplet formation during polymerization [11]. Calcium alginate gels are the most widely used biocompatible hydrogels due to their low cost, ease of use, and gentle gelation conditions.

However, alginate gels are limited by relatively poor mechanical strength and weak chemical stability under physiological conditions [9].

Since alginate is the most widely used biocompatible hydrogel material, a substantial body of literature has demonstrated scalable approaches for producing alginate gel particles across sub-millimeter size range [9, 12]. If an alginate-based gel formulation could be developed with enhanced chemical stability and mechanical robustness, it could serve as a practical platform for developing encapsulated microbes as an orally deliverable treatment for CKD.

Alginate can be enhanced through blending with other polymers such as PVA, polyacrylamide, chitosan, starch, besides others and it can be cross-linked by a range of divalent and trivalent cations that bind to alginate more strongly than does calcium [8, 10]. Barium ions are frequently used to improve alginate resistance to dissolution, while copper is known to produce strong alginate gels [13]. The trivalent cations such as aluminum and ferric iron display high affinity for alginate [14] and can yield substantially strong networks [15, 16]. Among these candidates, ferric iron is expected to be the least toxic, so we have pursued the development of ferric alginate gels as a promising path to a robust alginate formulation suitable for oral administration, intact passage through the gut, and practical production at scale. Together, these considerations motivate the development of ferric-crosslinked alginate hydrogels as a scalable, biocompatible, and mechanically robust platform for microbial encapsulation and oral delivery in CKD therapy.

## Results and Discussion

### Alginate crosslinked by ferric iron

There are reports in the literature of producing ferric alginate beads by dropping sodium alginate solution into an FeCl_3_ solution [17]. However, this is not likely to work for encapsulation of live cells due to the very acidic pH of FeCl_3_ solutions. Adjusting the pH of the FeCl_3_ solution, which initially has a pH of 1.5, to a value above 3 results in precipitation so we use ferric citrate solution, which can be adjusted to pH 6 without inducing immediate precipitation. Dropping a sodium alginate solution into a 50mM ferric citrate solution results in the formation of gel beads that have an opaque solid surface and still have a liquid core after 2 hours in the ferric solution. Dropping sodium alginate into 10mM or 1mM ferric citrate still makes liquid core beads due to the very high binding affinity of Fe^+3^ to alginate rapidly producing a densely cross-linked surface layer that substantially impedes further penetration of Fe^+3^ into the core. Reports in the literature indicate that a more homogeneous ferric alginate gel can be produced by first making calcium alginate and then exchanging some of the calcium ions for ferric ions or by creating a competition for ion binding sites by using an excess of Ca^+2^ over Fe^+3^ in the cross-linking bath [18, 19]. Using a 6-fold molar excess of sodium over calcium has been reported to result in a homogeneous distribution of calcium in alginate gels [20]. We have found that combining this with a 10-fold excess of Ca^+2^ over Fe^+3^ in the gelation bath allows production of homogeneous solid gel beads of yellow to brown color and which contains iron in addition to calcium.

### Mixed gel of polyvinyl alcohol and alginate crosslinked by ferric iron

The combination of 6% PVA with 2% sodium alginate (SA) crosslinked by calcium is more elastic and resilient than pure calcium alginate and it was used for in-vitro experiments demonstrating that the cresol-degrading bacterium *Thauera aminoaromatica* could be encapsulated in a hydrogel and used to degrade p-cresol at physiologically relevant concentrations [4]. The same 6%/2% PVA/SA gel cross-linked with Ba^+2^ was shown to be stable for 6 months in a bioreactor for removing nitrogen from simulated municipal wastewater [21] so we included 6%/2% PVA/SA gel crosslinked with Ca^+2^ and Fe^+3^ in our screen for gel stability under simulated gastrointestinal tract conditions.

### Interaction of various alginate-based gel compositions with low MW chitosan

It has been reported that chitosan of 20kDa MW or less can penetrate into calcium alginate gels and provide additional cross-linking [20, 22], which presumably occurs through formation of polyelectrolyte complexes (PECs) with the negatively charged alginate. Chitosan is also known to form very strong complexes with ferric ions [23] so it is possible that a ferric alginate gel could contain both PEC-related cross-linking and ferric complex related crosslinking when chitosan is present within the gel matrix. To assess whether these molecular interactions could enhance the stability of an alginate gel we began comparing candidate alginate formulations by performing chemical stability testing consisting of incubating gel beads in simulated gastric fluid at pH 3 for 4 hours at 37C with gentle shaking and then in simulated intestinal fluid at pH 6 for 16 hours at 37C with shaking, and by doing compression testing of gel beads using a texture analyzer. The results of these initial gel formulation screening tests are summarized in Table 1.

**Table 1.**
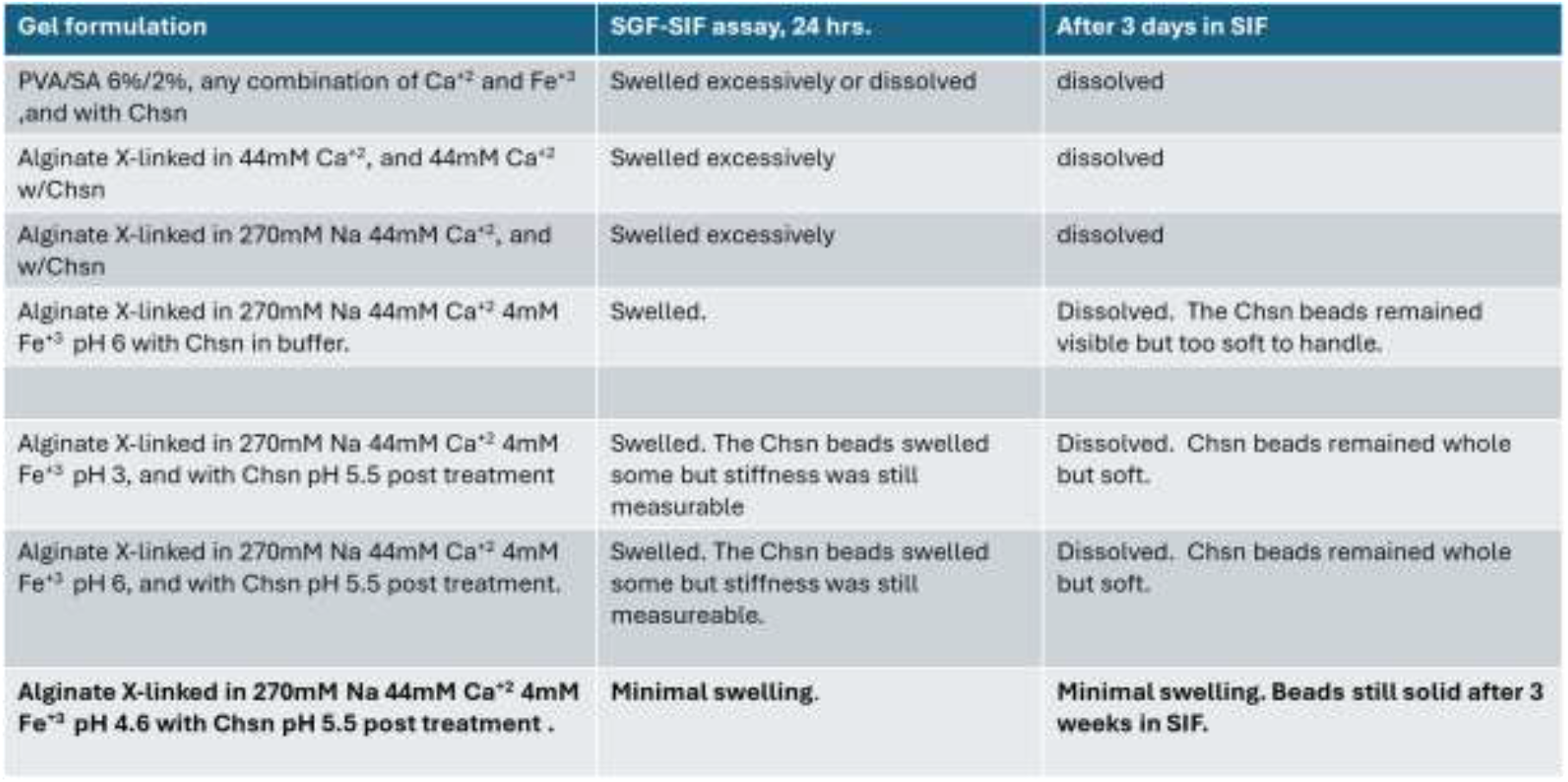
Summary of initial gel formulation screening results. Gel beads of about 2mm diameter were produced as described in Materials and Methods. Chitosan-treated beads were prepared by rinsing the cross-linked beads in water and then stirring them in 0.2% 20kDa chitosan solution. The gel beads were then subjected to the SGF-SIF assay to determine of the gel was stable enough to remain intact during a transit of the mammalian gastro-intestinal tract.

| Gel formulation | SGF-SIF assay, 24 hrs. | After 3 days in SIF |
| --- | --- | --- |
| PVA/SA 6%/2%, any combination of $\text{Ca}^{+2}$ and $\text{Fe}^{+3}$, and with Chsn | Swelled excessively or dissolved | dissolved |
| Alginate X-linked in 44mM $\text{Ca}^{+2}$, and 44mM $\text{Ca}^{+2}$ w/Chsn | Swelled excessively | dissolved |
| Alginate X-linked in 270mM Na 44mM $\text{Ca}^{+2}$, and w/Chsn | Swelled excessively | dissolved |
| Alginate X-linked in 270mM Na 44mM $\text{Ca}^{+2}$ 4mM $\text{Fe}^{+3}$ pH 6 with Chsn in buffer. | Swelled. | Dissolved. The Chsn beads remained visible but too soft to handle. |
| Alginate X-linked in 270mM Na 44mM $\text{Ca}^{+2}$ 4mM $\text{Fe}^{+3}$ pH 3, and with Chsn pH 5.5 post treatment | Swelled. The Chsn beads swelled some but stiffness was still measurable | Dissolved. Chsn beads remained whole but soft. |
| Alginate X-linked in 270mM Na 44mM $\text{Ca}^{+2}$ 4mM $\text{Fe}^{+3}$ pH 6, and with Chsn pH 5.5 post treatment. | Swelled. The Chsn beads swelled some but stiffness was still measurable. | Dissolved. Chsn beads remained whole but soft. |
| Alginate X-linked in 270mM Na 44mM $\text{Ca}^{+2}$ 4mM $\text{Fe}^{+3}$ pH 4.6 with Chsn pH 5.5 post treatment . | Minimal swelling. | Minimal swelling. Beads still solid after 3 weeks in SIF. |

The stiffness of the alginate beads prepared using the 3 ferric citrate-containing gelation baths with different pH’s were measured on a texture analyzer for comparison. The beads produced in the pH 3 bath were the stiffest by a large margin, but these were not the most stable beads in the SGF-SIF stability assay. A gelation bath with a pH of 3 is also likely to reduce cell viability for at least some bacteria that would be useful for metabolizing uremic toxin precursors. Treating calcium-ferric alginate (Ca-Fe^+3^-alginate) beads with chitosan improved their stability in the SGF-SIF assay even though in the case of the pH 3 and pH 6 baths it resulted in a reduction of bead stiffness. The Ca-Fe^+3^ alginate beads prepared at pH 4.6 and then treated with chitosan were both the most stable in the SGF-SIF assay and the only ones which became stiffer after the chitosan treatment, Figure 1.

**Figure 1.**
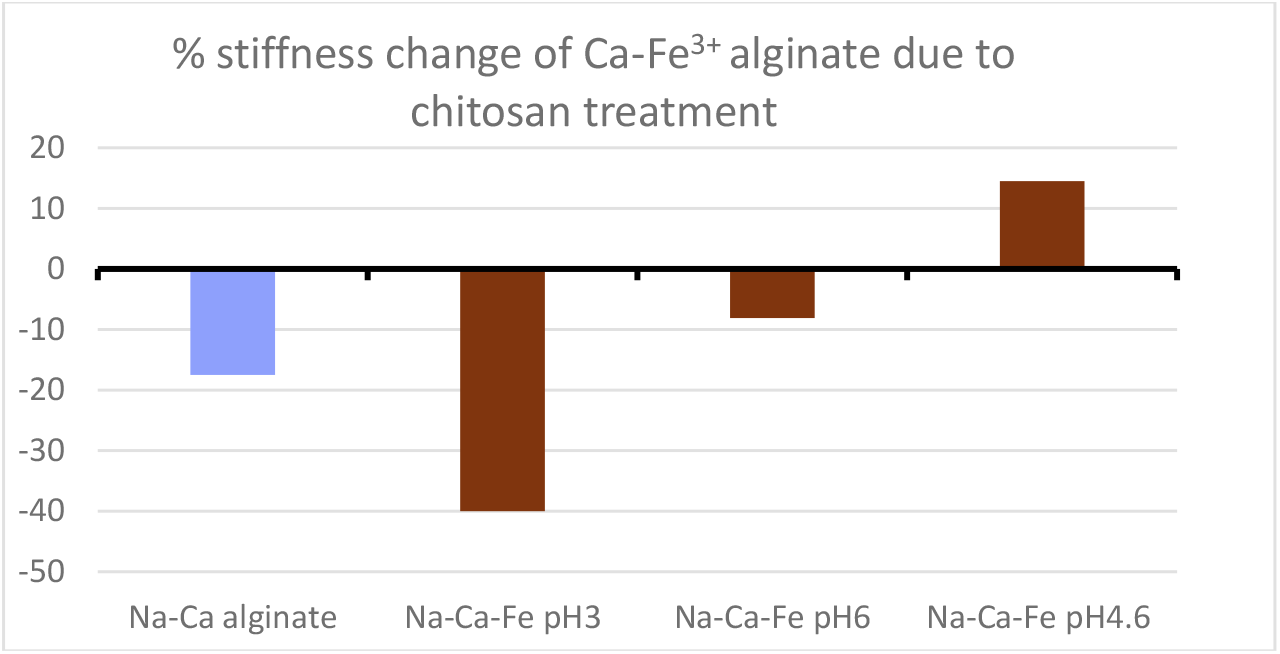
Effect of chitosan treatment on the stiffness of 4 ferric iron-containing alginate gel bead formulations.

### Cell viability optimization and p-cresol degrading activity in Ca-Fe^+3^-alginate chitosan-treated beads

The chitosan-treated Ca-Fe^+3^ pH 4.6 alginate beads meet the basic requirements to be used for small animal testing so we attempted encapsulation of *Thauera aminoaromatica* cultures in this gel using different amounts of time in the gelation bath and subsequently in the chitosan bath to optimize the gelation protocol. *T. aminoaromatica* grows well anaerobically using acetate as carbon source and nitrate as electron acceptor so conversion of nitrate to nitrite in acetate medium was used as an assay for cell viability, Figure 2. The viability of encapsulated *T. aminoaromatica* drops dramatically between 2 and 4 hours in the Na270Ca44Fe4 pH 4.6 gelation solution. Four hours in the 0.2% chitosan solution was chosen as the standard protocol since it allows the gel beads to be conveniently produced in one day. As shown in Figure 2, the cresol-metabolising activity of *T. aminoaromatica* encapsulated in the Ca-Fe^+3^ pH 4.6 alginate gel beads lasts for at least 7 days.

**Figure 2.**
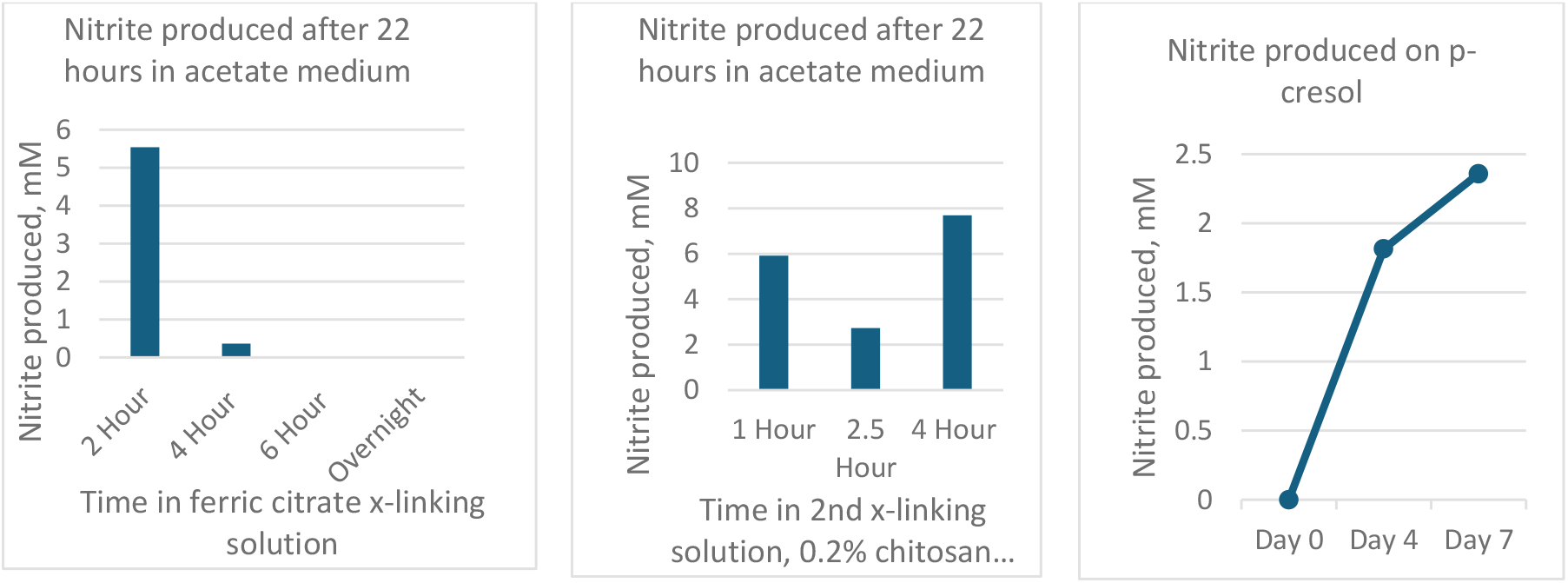
Optimizing the encapsulation process for *Thauera aminoaromatica* in 2mm diameter Ca-Fe^+3^ pH 4.6 2% alginate beads. Left panel – gel beads should not remain in the pH 4.6 gelation bath for longer than 2 hours. Center panel – time spent in the pH 5.5-6 chitosan bath makes little difference to cell viability. Right panel – Confirming that *T. aminoaromatica* encapsulated using the 2hr/4hr gelation protocol is capable of metabolizing p-cresol.

### Cell viability and p-cresol degrading activity in 400 micron Ca-Fe^+3^-alginate chitosan-treated beads

To confirm that active *T. aminoaromatica* beads could be produced in a size and quantity suitable for rodent testing using the 2hr/4hr gelation protocol for the Ca-Fe^+3^ pH 4.6 alginate gel we prepared a 50mL batch of 400 micron 1% alginate beads using a Buchi encapsulator configured with a 200 micron dropping orifice. After completion of the 2hr/4hr gelation protocol the beads were collected in a sanitized 250 micron sieve, rinsed with MilliQ water, placed in acetate growth medium, and incubated anaerobically at 37°C overnight. After moving the beads into p-cresol-containing anaerobic media 0.25mM p-cresol was fully metabolized within 22 hours, Figure 3.

**Figure 3.**
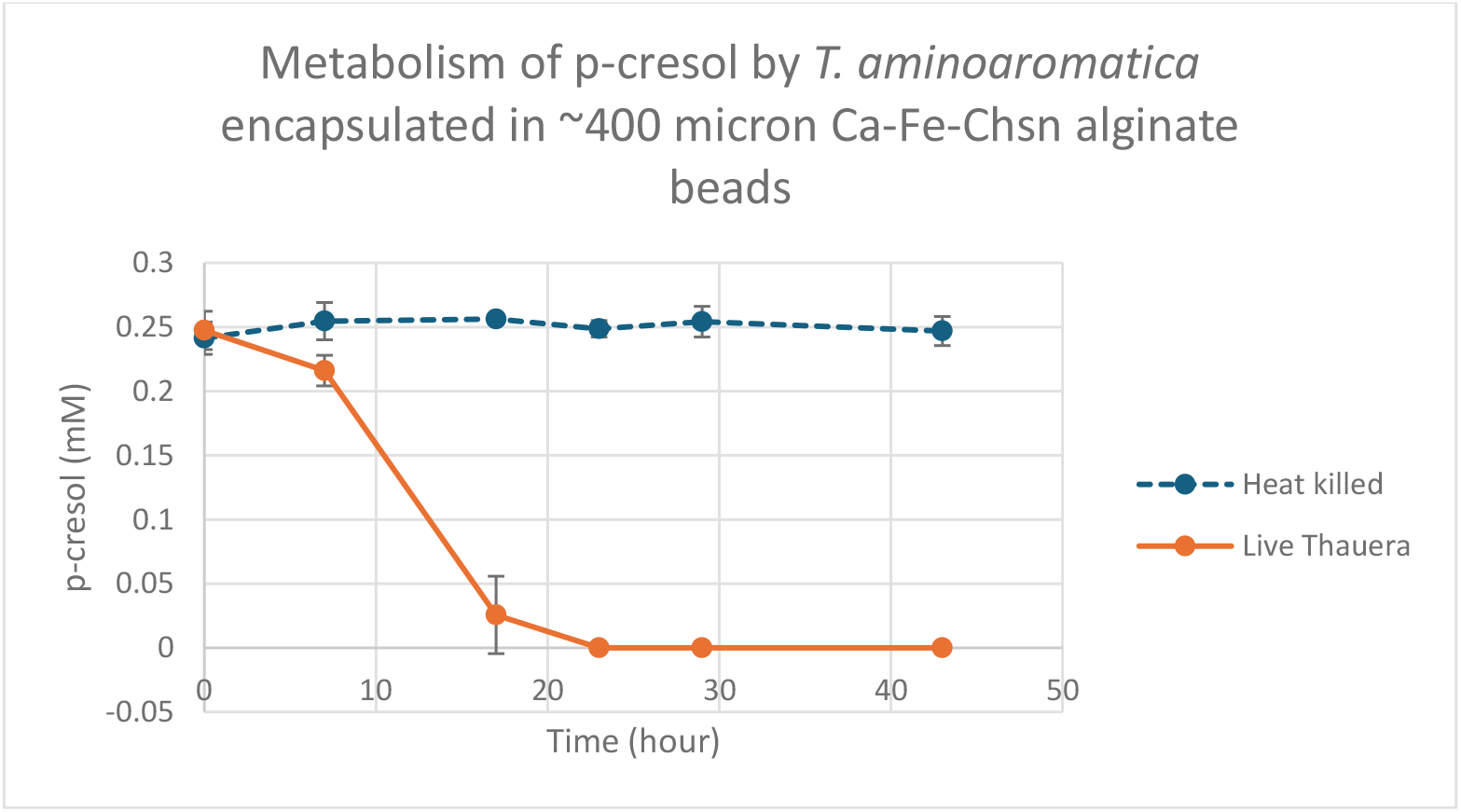
Live *T. aminoaromatica* cells and heat-killed *T. aminoaromatica* cells encapsulated in 400 micron Ca-Fe^+3^-alginate-chitosan beads incubated in MRS medium containing 0.25mM p-cresol for 2 days. Concentration of p-cresol was measured by HPLC assay at indicated time points. The p-cresol was fully removed within 24 hours, confirming the suitability of the encapsulation method for use in rodent trials.

### Analyzing the contribution of pH to the stabilization of this alginate gel by ferric iron and chitosan

The pH of the gelation solution used for the initial formation of alginate gel beads was found to be an important parameter affecting the stiffness and chemical stability of the final chitosan-treated beads. Unadjusted solutions containing ferric citrate have pH of about 3. Unadjusted solutions containing CaCl_2_ have a pH of about 6, the condition used in the majority of the literature concerning calcium alginate gels. A pH of 3 may not be practical for encapsulating a wide range of bacteria so we tried to determine the highest pH at which calcium ferric alginate gel beads could be produced and then be stabilized when treated with chitosan. We made stiffness measurements on simple Ca alginate gel beads prepared at pH 4.5 and pH 6, and on homogeneous calcium alginate gel beads produced with a 6-fold excess of NaCl to CaCl_2_, each prepared at pH 4.5 and pH 6 to try to determine if the gelation pH alone is a strong contributor to the stabilization observed in the calcium ferric alginate gel treated with chitosan. In the case of the simple calcium alginate gel we found that stiffness of gel beads produced at pH is substantially lower than those produced at pH 6, and in both cases treatment with chitosan significantly reduced the measured stiffness. In the case of the homogeneous gel beads produced with a 6-fold excess of NaCl to CaCl_2_, the pH has no effect on the stiffness with or without chitosan treatment, Figure 4a., even though the homogeneous alginate gels absorb and incorporate a substantial amount of chitosan, Figure 4b. These results suggest that the interaction between chitosan and the ferric iron in the Na270Ca44Fe4 crosslinked gels is the primary contributor to the stabilization of the Ca-Fe^+3^ pH 4.6 alginate gel formulation.

**Figure 4.**
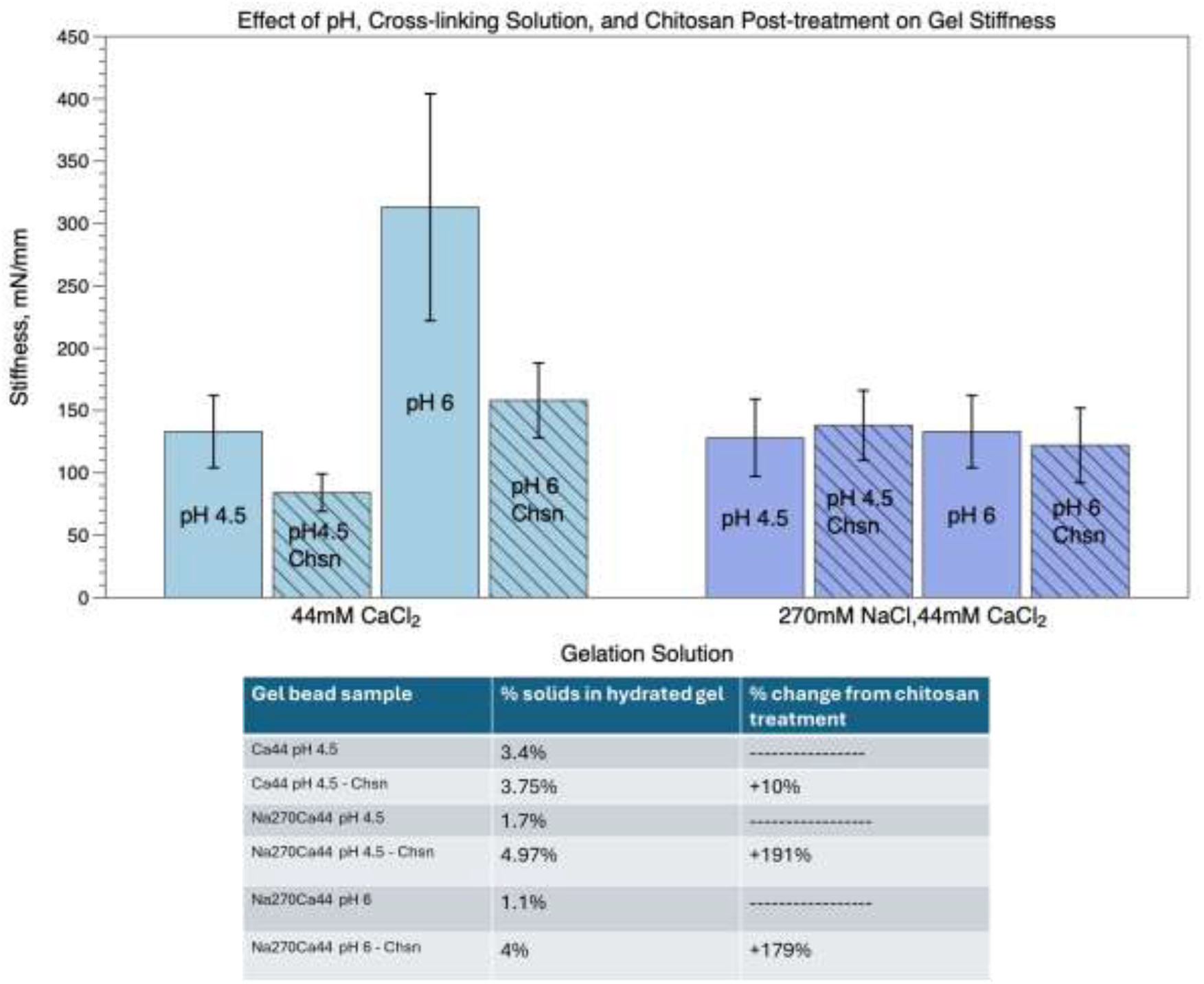
– Top panel - Gel beads were made by dropping 1% sodium alginate solution into either 44mM CaCl_2_ or 270mM NaCl, 44mM CaCl_2_ at the two pHs indicated. Stiffness values from 5 sequential measurements on 10 beads of each type are shown. Bottom panel - Change in solids content of the gels after chitosan treatment.

### Producing the Ca-Fe^+3^ pH 4.6 alginate gel in slab form for physical measurements

In order to obtain accurate measurements of the elastic modulus of the gel and how it is affected by the presence of ferric ions, chitosan, and both combined, as well as to be able to obtain IR spectra, we needed to produce flat gel slabs. This was done by preparing agarose molds as described in Materials and Methods. The gel slabs produced were about 1.8mm thick, 10cm long, and 5 cm wide. After equilibrating each gel sample in several changes of water over 48 hours 1cm x 3cm strips were cut from each gel slab and used for collecting stress/strain curves using a 2mm^2^ compression probe at 10 different locations dispersed over the area of the sample, Figure 5b. Wet and dry mass measurements were also made on duplicate samples of each gel, Figure 5a.

**Figure 5.**
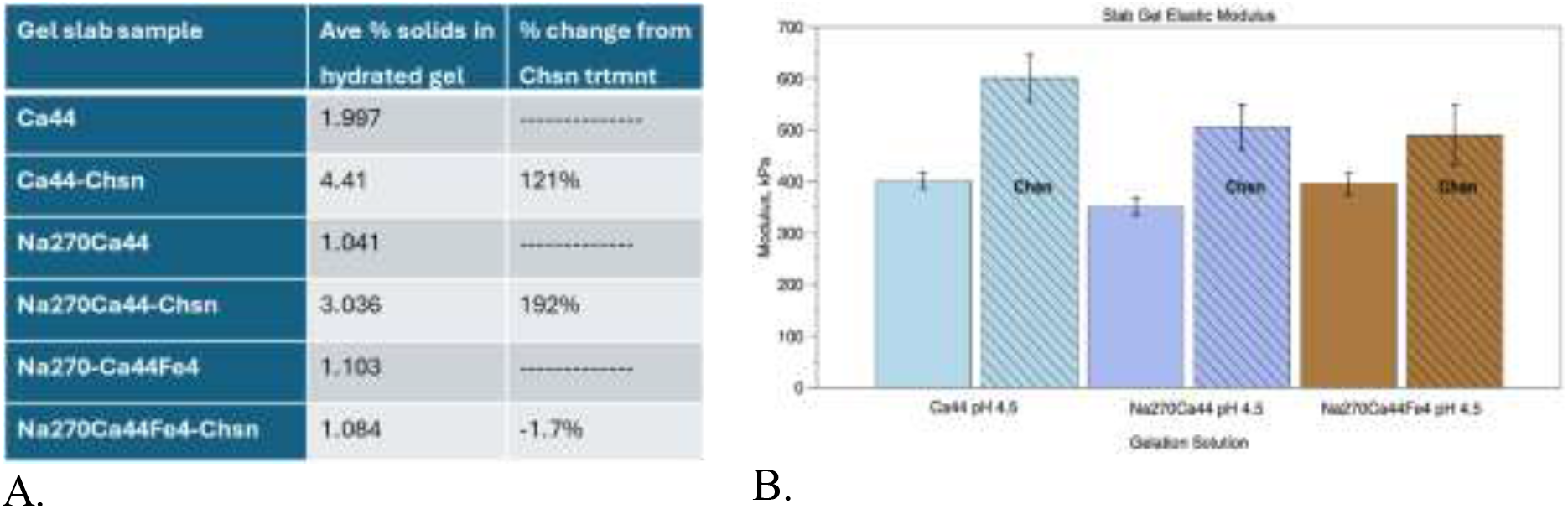
a - Change in solids content of the gels after chitosan treatment. Figure 5b – Elastic modulus measurements on the gels.

As was seen in the gel beads, chitosan is absorbed into the Na270Ca44 gels, however, minimal uptake of chitosan was seen in the Ca44 gel beads whereas the dry mass of the Ca44 gel slab more than doubled during the chitosan treatment. This variation may be due to differences in Ca^+2^ diffusion kinetics during gelation in a stirred solution as compared to diffusion through a static agarose gel. An important observation from the slab gel experiment is that even though there is an increase in the modulus of the chitosan treated Na270Ca44Fe4 gel slab there is no measurable uptake of chitosan into this gel. This suggests that the modulus change is due to chitosan absorption onto the surface of the gel. The very strong binding of ferric iron to chitosan is apparently inhibiting the penetration of chitosan into the iron-containing alginate gel matrix.

### IR spectroscopy

Pieces of the gel slabs were allowed to dry at room temperature in ambient air for 1 week. FT-IR spectra were collected using an ATR cell from two to four points on each sample and were found to be indistinguishable. Figure 6 shows the calcium alginate gel spectra overlaid with the chitosan-treated gel spectra. Clear differences are seen between the plain and chitosan-treated samples of the Ca44 gel slabs and the Na270Ca44 gel slabs in the main asymmetric and symmetric and carboxyl stretch peaks at 1589cm^-1^ and 1418cm^-1^. The presence of chitosan in the Ca44 and Na270Ca44 gels causes a reduction in height of the carboxyl peaks. This is likely due to the formation of a charge complex which may be partially damping the carboxyl peaks in these samples, Figure 5a.

**Figure 6.**
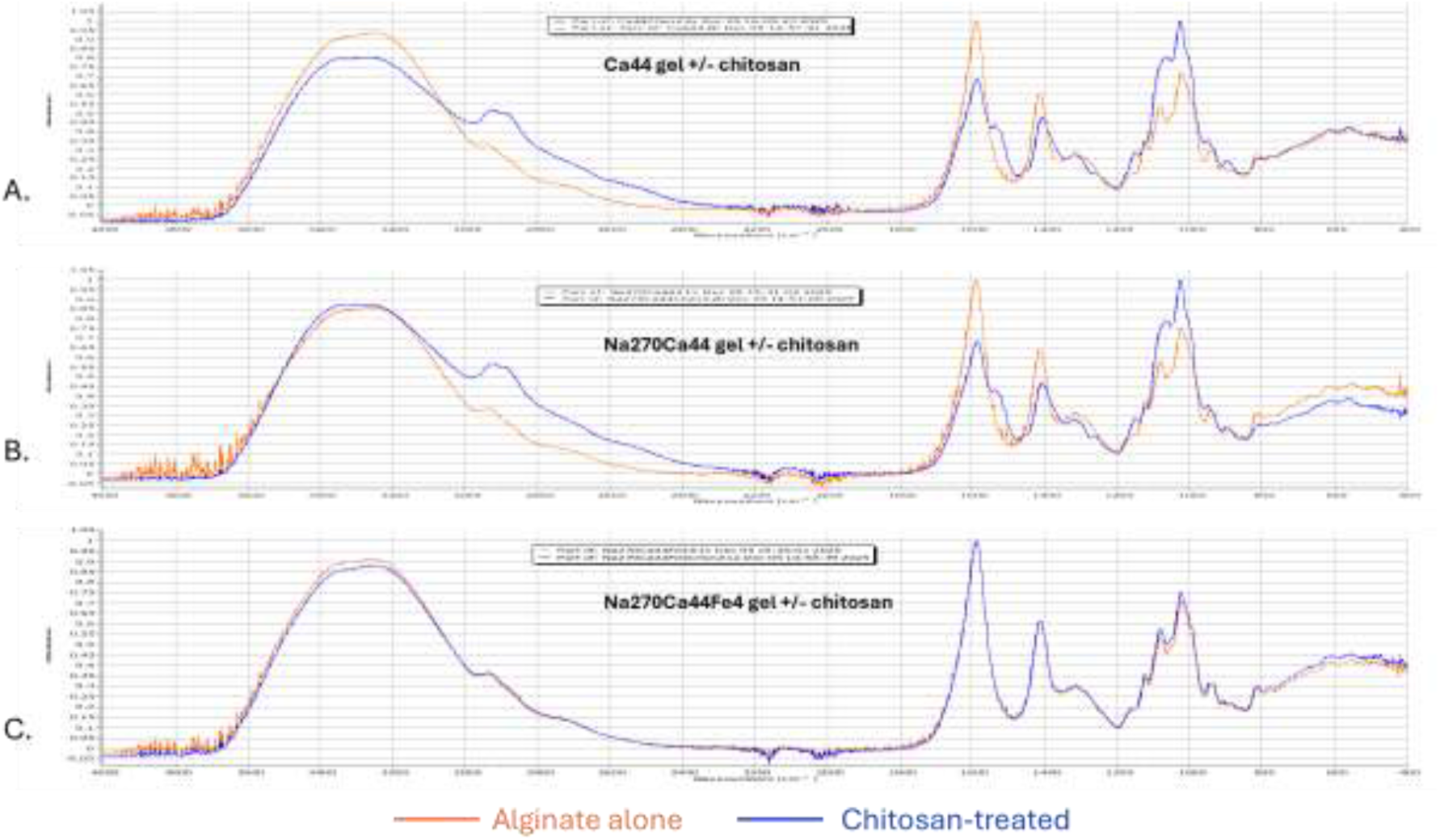
FT-IR spectra of the 3 types of slab gels with and without chitosan treatment. Panel A - Ca44 calcium alginate gel with and without chitosan treatment. Panel B - Na270Ca44 homogeneous calcium alginate gel with and without chitosan treatment. Panel C – Na270Ca44Fe4 calcium ferric alginate gel with and without chitosan treatment.

No difference is seen in the carboxyl stretch peaks or elsewhere in the spectrum for the Na270Ca44Fe4 gel slabs with and without chitosan treatment, confirming the lack of chitosan absorption into the ferric iron-containing alginate gel. The presence of ferric ions in the gel did not result in an obvious shift in position of the carboxyl bands. This may reflect a relatively low occupancy of iron in the di/tri-valent ion binding sites.

## Conclusions

In this study, we demonstrate that a controlled incorporation of ferric iron into calcium alginate hydrogels can be achieved through a three-ion competition strategy, enabling the formation of ferric-containing calcium alginate gels without substantially altering the bulk mechanical properties relative to conventional calcium alginate. The physical properties of the resulting ferric-containing calcium alginate gel were not substantially different from a simple homogeneous calcium alginate gel. However, after treatment of the ferric-containing calcium alginate gel that was formed at pH 4.6 with a chitosan solution adjusted to a pH of 5.5-6 a change occurs which dramatically increases the stability of the gel to physiologic salt concentrations at acidic and neutral pHs. The ferric-containing calcium alginate gel that was formed at pH 4.6 does not absorb a measurable amount of chitosan while similar calcium alginate gels which do not contain iron do absorb a substantial amount of chitosan under the same conditions. Therefore we conclude that the change in properties we observed in the ferric-containing calcium alginate relative to calcium alginate gels is due to the formation of a thin layer of chitosan complexing with ferric ions and alginate at the surface of the gel and that these complexes become dense enough to block further penetration of chitosan into the bulk of the gel matrix.

Ferric ions have been shown to bind to both MG blocks and GG blocks in alginate [17], whereas calcium binds primarily at the GG blocks as depicted in the eggbox model [24]. It is possible that in a mixed calcium and ferric iron alginate gel produced with an excess of calcium most of the GG blocks are occupied by calcium and some fractions of the MG blocks are occupied by ferric ions. This may explain why this Ca-Fe+3 gel has properties similar to those of a homogenous calcium alginate gel. Since ferric ions in alginate complexes are in a hexacoordinate state it would be possible for chitosan to form bridges between sites where iron is bound to two MG blocks [23]. A model for this structure is proposed in Figure 7. Our present data does not explain why the CaFe^+3^ alginate gel produced at pH 4.6 behaves differently when treated with chitosan than the same gel produced at pH6 or pH3. This pH effect may be attributed to the degree of hydrolysis of ferric ions in solution as is suggested by the colloidal model of interaction rather than to the structure of the alginate gel formed at this pH [25].

**Figure 7.**
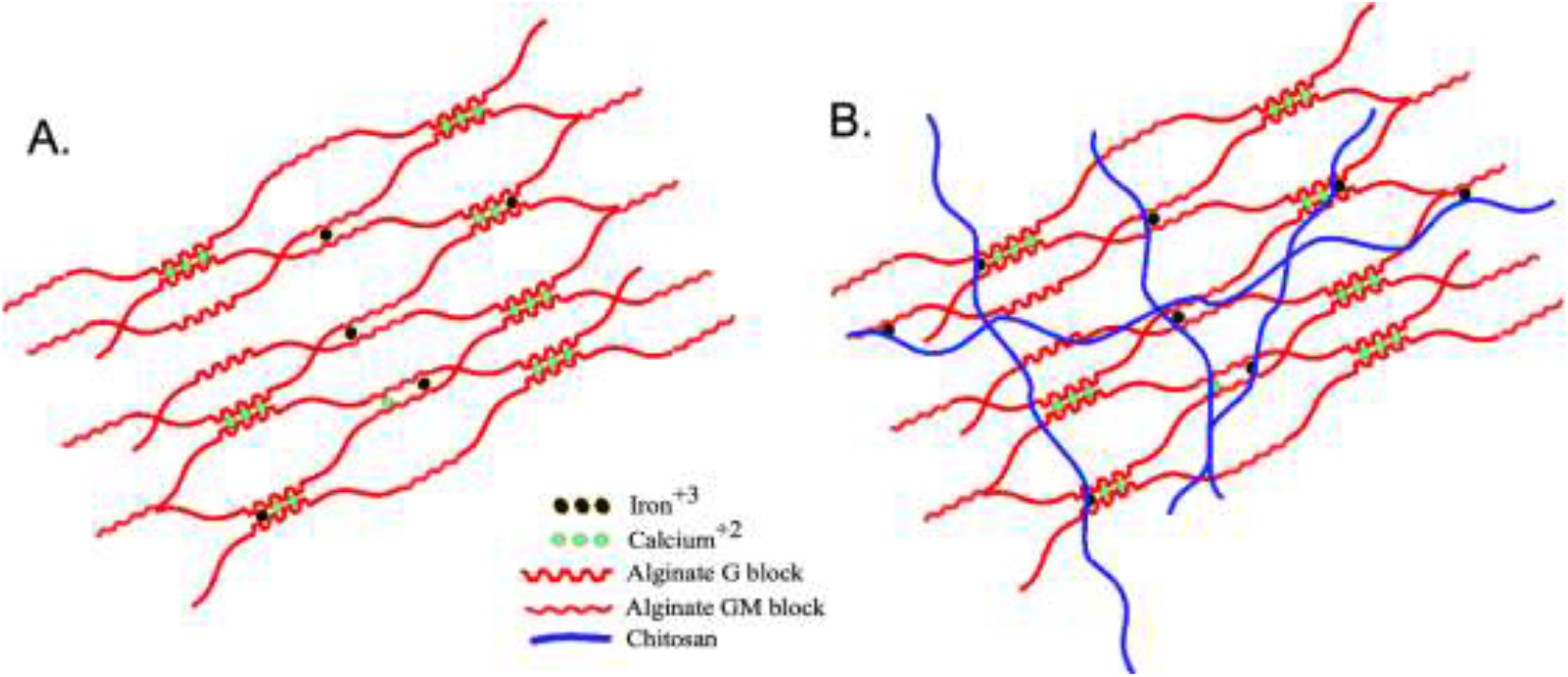
A. Proposed molecular network of the ferric-modified homogenous calcium alginate gel. B. Surface layer of the network as modified by chitosan binding to available ferric ions.

Even though the thin layer on the surface of the CaFe^+3^ alginate gel that forms after addition of chitosan does not contribute measurable mass to the gel it is robust enough that it increases both the modulus of the gel and prevents the dissolution of the underlying alginate matrix due to salt and pH exposures typically found in the gastro-intestinal tract of mammals. This hybrid alginate gel can be used to produce sub-millimeter gel beads which can encapsulate live *Thauera aminoaromatica* culture [4], maintaining it’s ability to degrade p-cresol under anaerobic conditions at 37C using nitrate as electron acceptor. Collectively, these findings establish ferric-modified alginate gels as a promising and scalable platform for robust oral delivery of encapsulated microbial therapeutics.

## Materials and Methods

Sodium alginate from brown algae was obtained from Sigma-Aldrich, St Louis MO. Chitosan with a MW of 20 kDa was obtained from Glentham Life Sciences, Corsham, UK. All other chemicals used were of reagent grade and supplied by Fisher Scientific, USA or VWR Inc., USA.

### Initial screening of gel beads prepared using different gelation solutions

Small batches of millimeter sized calcium alginate and polyvinyl alcohol alginate (PVA/SA) gel beads used during the initial screening of gel formulations were made by dropping 2% sodium alginate (SA) solution from a 10 mL syringe fitted with a 26 ga needle into a hardening bath consisting of 44mM CaCl_2_ (Ca44), 270mM NaCl-44mMCaCl_2_ (Na270Ca44), or 270mM NaCl-44mMCaCl_2_-4mMFe^3+^ citrate (Na270Ca44Fe4). These solutions were either used as made or with pH adjusted to values given in the Results and Discussion section. The gel beads were left stirring in the gelation solution for 2 to 4 hours before being collected in a sieve and rinsed with water. The beads being treated with chitosan were subsequently placed in 10 volumes of a 0.2% solution of 20kDa chitosan adjusted to a pH of 5.5 to 6 using pH paper and left to stir slowly overnight. The untreated beads were placed in 10 volumes of water and left stirring overnight. The following day both the chitosan-treated and untreated beads were placed in fresh water which was changed 3 times over a period of 4 hours before stiffness measurement were made using a Texture Analyzer or samples were subjected to the simulated gastric fluid (SGF) and simulated intestinal fluid (SIF) assay.

### Simulated gastric fluid-simulated intestinal fluid (SGF-SIF) stability assay

MRS media as defined by DSMZ, Leibnitz Institute, Braunschweig-Süd, Germany, was used as the basis of SGF and SIF buffers. The composition of MRS medium is as follows, values are per liter:

Bacto proteose peptone: 5 g, Beef extract: 5 g, Yeast extract: 5 g, Glucose: 10 g, Tween 80: 1 ml, K2HPO4: 0.2 g, Na-acetate: 5 g, (NH4)3 citrate: 2.4 g, MgSO4X7H2O: 0.2 g, MnSO4XH2O:0.05 g, NH4Cl: 0.2 g

For SGF the pH is adjusted to 3.2. For SIF the pH is adjusted to 6.5. The SGF-SIF assay is conducted by placing 2 gm of gel beads into 10mL of SGF in a Balch tube and shaking slowly in the slanted tube for 2 hours at 37C. Then the SGF is withdrawn, beads are rinsed with 5mL of deionized water once, and then the water is replaced by 10mL of SIF. The beads then shake gently overnight (∼16hrs.) at 37C.

### Texture Analysis

Stiffness measurements of individual gel beads were made using a TA.XT Express Connect, Stable Microsystems, Surrey, UK, equipped with a 5 Kg load cell and a 10mm cylindrical plastic probe with flat end. Gel beads were placed on a glass microscope slide in a drop of water. A return-to-start cycle is used where the probe is lowered until the contact trigger force of 100mg is detected, at which point the compression continues until 85% of the height at trigger force is reached and then the probe is withdrawn. This cycle is repeated 5 times, recording a force-distance curve for a 15% compression of the bead each cycle. The data for the 5 cycles are averaged. The 5 cycle averages for 10 beads are reported as the stiffness for each gel bead formulation.

For elastic modulus measurements on flat gel slab samples a teflon probe with a 2mm^2^ area was slowly lowered onto a roughly 1cm x 3cm flat gel sample kept wet with a film of water pipetted onto the surface. The height at the point of trigger force detection is taken as the strain height and 5 stress measurements to 15% of strain height are made at each of 10 points distributed over the surface of the sample. The 5 stress-strain curves are averaged for each point. The average of the slopes of the stress-strain curves at the 10 measurement points is reported as the elastic modulus in kPa for each gel formulation.

### *Thauera aminoaromatica* cultures for encapsulation

Growth of *Thauera aminoaromatica* cultures was done as described (Prakit’s paper). Briefly, a 10mL culture of *T. aminoaromatica* was grown under nitrogen in a Balch tube to an OD of 1.2 over 72 hours in UGA medium containing 32mM acetate as carbon source and 10mM nitrate as electron acceptor. The 10mL culture was used to inoculate 150 mL of the same medium in a serum bottle and was grown for 48 hours under nitrogen. The cells were collected by centrifugation and then resuspended in 150mL of fresh UGA containing 0.6mM p-cresol and 10mM nitrate as carbon source and electron acceptor and incubated overnight at 37C to induce expression of the p-cresol utilization enzymes. The cells were then collected by centrifugation and resuspended in 4mL of UGA media with acetate and nitrate.

A 2.5% stock sodium alginate (SA) solution was prepared by heating the SA powder in MilliQ water to boiling 3 times in a microwave oven with intermittent stirring. 2.3mL of the cell concentrate was mixed with 20mL of 2.5% SA 15mL of growth medium and 24mL of MilliQ water in a sterile 50mL Falcon tube to produce 50mL of 1% SA solution containing live *T. aminoaromatica* cells. A smaller batch of heat killed cells were prepared to make negative control gel beads. 0.67mL of the cell concentrate was placed in an 85C water bath for 30 minutes and then mixed with 2.5% SA and water to make 20mL of 1% SA containing heat-killed cells.

### Cell Encapsulation

Sub-millimeter alginate gel beads used for encapsulating live cell cultures of were produced using a Buchi B390 Encapsulator, an instrument which uses acoustic waves to break up a fine jet of liquid sodium alginate solution into a train of small droplets. The size of the droplets produced are a function of the orifice diameter producing the liquid stream and the frequency of the acoustic energy fed into the liquid immediately upstream of the orifice. The feed container and tubing for the Encapsulator and a 10L tank to hold the gelation solution and mechanical stirrer were sanitized using household bleach and rinsed with MilliQ water. The Buchi Encapsulator head and an inline 60 micron nylon screen filter were soaked in 70% ethanol for 20 minutes then rinsed with MilliQ water.

First the 20mL of 1% SA containing the heat killed cells was placed in the Encapsulator where it passed through a 200 micron orifice and was dispersed as a stream of uniform small droplets into 3L of gently stirred Na270Ca44Fe4 pH 4.6 gelation solution in about 10 minutes. Then the gelation solution containing the beads was transferred into a 4L beaker and stirred slowly for 2 hours. 4L of fresh gelation solution was placed in the tank and the 50mL of 1% SA containing live *T*.*a*. cells was dropped in as above over about 20 minutes and allowed to continue stirring for 2 hours.

After the two batches of roughly 400 micron beads were stirred gently for 2 hours in the gelation solutions they were collected in a 250 micron sanitized sieve, rinsed with MilliQ water, and placed into 10 volumes of 0.2 micron filtered 0.2% chitosan solution as described above and stirred gently for 4 hours. The two batches of gel beads were collected in sanitized sieves, rinsed with MilliQ water, and placed in serum bottles containing 250 mL of UGA medium with acetate as carbon source [4] and flushed with nitrogen gas for 20 minutes. The anaerobic live and control encapsulated cultures were incubated overnight at 37C.

### Cresol degradation activity test

Three grams of gel beads containing live or heat-killed *T. aminoaromatica* were placed in serum bottles with 20mL of UGA media containing 32mM acetate as carbon source and 10mM nitrate as electron acceptor, flushed with nitrogen, and incubated at 30C overnight with shaking to allow metabolic activity to recover from the encapsulation process. Subsequently the acetate medium was removed and replaced with 40mL of UGA medium containing 0.25mM p-cresol as carbon source and 10mM nitrate as electron acceptor, flushed with nitrogen, and maintained under anaerobic conditions with shaking at 37C. Samples of culture fluid were collected at the indicated time points and analyzed by HPLC as described [4].

### Production of slab gels

Reusable mold blanks were fabricated from two layers of 1/16” polycarbonate sheet in the shape of tapered rectangles. The blanks are about 1.8mm thick and the taper helps them release suction as they are pulled out of the cooled and hardened cast agarose, leaving behind a thin slot in the cast agarose which can be filled with SA solution. The gelation salt solutions are made at 1.2x concentration and the pH is adjusted to a value about 0.6 – 0.7 pH units lower than the target value of pH 4.5. Agarose is dissolved in deionized water at 4% and then mixed while still hot with the pre-warmed 1.2x gelation salt solutions to produce 0.8% agarose containing 1x gelation buffers. The pH of the warm agarose solutions are then checked and adjusted to pH 4.5 using a mixture of bromocresol green and methyl red dyes which turns grey at this pH value, Fisher Scientific cat. R121500060A. 210 mL of the warm pH-adjusted agarose solution is then poured into a rectangular plastic container and the 1.8mm thick mold blanks are inserted to full depth and clamped in position while the agarose cools and hardens over night. The next day the mold blanks were carefully removed from the agarose and the slot was filled with 9 mL of the same pH 4.5 gelation solution present in the cast agarose. After a few hours the pH of the fill solution is checked using a pH meter to verify whether or not the target pH has remained stable. If the pH has drifted the solution is drained and replaced with fresh pH 4.5 solution and the process is repeated. This required a number of cycles over several days in the case of the ferric citrate-containing gelation buffer as agarose seems to interfere with the ferric iron in this solution.

Once stable pH values were reached the gel slots were filled with water for 2 minutes and then emptied, repeated twice. Immediately after the second deionized water was drained out the 1% SA solution was introduced into the slot from a syringe through a long 18ga blunted needle starting from the bottom and continuing upwards as the needle was withdrawn. The alginate was left to gel in the molds for about 2 hours, long enough for the gel to become solid enough to handle, and then the soft alginate slabs are pulled out of the agarose molds and placed into a container of the gelation buffer for another 2 hours to allow gelation to finish under free solution conditions. The gel slabs were then rinsed in deionized water and cut in half. One half was placed in 0.2% chitosan solution for 4 hours and the other half was placed in water. Both halves were then soaked in several changes of water over the next 24 hours to ensure the gels were fully equilibrated in water before elastic modulus measurements were made.

### IR spectra

Sample preparation for collecting FT-IR spectra using ATR mode consisted of simply allowing small pieces of the gel slabs to dry in ambient indoor air for a week. This gentle drying would be expected to allow the most tightly bound fraction of the original water to remain in the gel structure. For comparison 5mL of the chitosan and SA solutions were placed in small petri dishes and allowed to dry down into thin sheets in order to collect IR spectra of these two pure components. A sample of dry alginic acid was prepared by soaking a piece of the dried sodium alginate sheet in 1N HCl, changing the HCl solution several times over 48 hours, and then soaking the hydrated alginic acid in deionized water with several changes over another 48 hours. The alginic acid was then allowed to dry into a thin solid sheet. Spectra were collected over wavelengths from 400/cm to 4000/cm with averaging over 64 reads on a Nicolet iS20 spectrometer using a diamond ATR cell.

## Acknowledgments

This project was funded in part by NIH 1R01DK130815-01 from NIDDK.

## References

1. Gryp, T., et al. Isolation and Quantification of Uremic Toxin Precursor-Generating Gut Bacteria in Chronic Kidney Disease Patients. International Journal of Molecular Sciences, 2020. 21, 1986 DOI: 10.3390/ijms21061986.

2. Jourde-Chiche, N. and S. Burtey, Accumulation of protein-bound uremic toxins: the kidney remains the leading culprit in the gut-liver-kidney axis. Kidney International, 2020. 97(6): p. 1102–1104.

3. Lin, C.-J., et al., Meta-Analysis of the Associations of p-Cresyl Sulfate (PCS) and Indoxyl Sulfate (IS) with Cardiovascular Events and All-Cause Mortality in Patients with Chronic Renal Failure. PLOS ONE, 2015. 10(7): p. e0132589.

4. Saingam, P., et al., Toward an effective delivery system of a microbial sink of the uremic toxin, p-cresol; an in vitro study with Thauera aminoaromatica S2. Frontiers in Microbiology, 2025. Volume 16 - 2025.

5. Candry, P., et al., Tailoring polyvinyl alcohol-sodium alginate (PVA-SA) hydrogel beads by controlling crosslinking pH and time. Scientific Reports, 2022. 12(1): p. 20822.

6. Banerjee, A., et al., Role of nanoparticle size, shape and surface chemistry in oral drug delivery. Journal of Controlled Release, 2016. 238: p. 176–185.

7. Kim, G.-H., Fluid and Electrolyte Problems in Chronic Kidney Disease, in Management of Chronic Kidney Disease: A Clinician’s Guide, M. Arıcı, Editor. 2023, Springer International Publishing: Cham. p. 327–344.

8. Hoffman, A.S., Hydrogels for biomedical applications. Advanced Drug Delivery Reviews, 2012. 64: p. 18–23.

9. Drury, J.L. and D.J. Mooney, Hydrogels for tissue engineering: scaffold design variables and applications. Biomaterials, 2003. 24(24): p. 4337–51.

10. Peppas, N.A., et al., Hydrogels in pharmaceutical formulations. Eur J Pharm Biopharm, 2000. 50(1): p. 27–46.

11. Nguyen, K.T. and J.L. West, Photopolymerizable hydrogels for tissue engineering applications. Biomaterials, 2002. 23(22): p. 4307–4314.

12. Lee, K.Y. and D.J. Mooney, Alginate: properties and biomedical applications. Prog Polym Sci, 2012. 37(1): p. 106–126.

13. Smidsrød, O. and G. Skjåk-Braek, Alginate as immobilization matrix for cells. Trends Biotechnol, 1990. 8(3): p. 71–8.

14. Tordi, P., et al., Cation-Alginate Complexes and Their Hydrogels: A Powerful Toolkit for the Development of Next-Generation Sustainable Functional Materials. Advanced Functional Materials, 2025. 35(9): p. 2416390.

15. Grant, G.T., et al., Biological interactions between polysaccharides and divalent cations: The egg-box model. FEBS Letters, 1973. 32(1): p. 195–198.

16. Malektaj, H., A.D. Drozdov, and J. de Claville Christiansen Mechanical Properties of Alginate Hydrogels Cross-Linked with Multivalent Cations. Polymers, 2023. 15, 3012 DOI: 10.3390/polym15143012.

17. Massana Roquero, D., et al., Iron(iii)-cross-linked alginate hydrogels: a critical review. Materials Advances, 2022. 3(4): p. 1849–1873.

18. Min, J.H. and J.G. Hering, Arsenate sorption by Fe(III)-doped alginate gels. Water Research, 1998. 32(5): p. 1544–1552.

19. Churio, O., F. Pizarro, and C. Valenzuela, Preparation and characterization of iron-alginate beads with some types of iron used in supplementation and fortification strategies. Food Hydrocolloids, 2018. 74: p. 1–10.

20. Klinkenberg, G., et al., Cell Release from Alginate Immobilized Lactococcus lactis ssp. lactis in Chitosan and Alginate Coated Beads. Journal of Dairy Science, 2001. 84(5): p. 1118–1127.

21. Godfrey, B., et al., Co-immobilization of AOA strains with anammox bacteria in three different synthetic bio-granules maintained under two substrate-level conditions. Chemosphere, 2023. 342: p. 140192.

22. Kopplin, G., et al., Alginate gels crosslinked with chitosan oligomers – a systematic investigation into alginate block structure and chitosan oligomer interaction. RSC Advances, 2021. 11(23): p. 13780–13798.

23. Hernández, R.B., et al., Coordination study of chitosan and Fe3+. Journal of Molecular Structure, 2008. 877(1): p. 89–99.

24. Donati, I. and B.E. Christensen, Alginate-metal cation interactions: Macromolecular approach. Carbohydrate Polymers, 2023. 321: p. 121280.

25. Sipos, P., et al., Formation of spherical iron(III) oxyhydroxide nanoparticles sterically stabilized by chitosan in aqueous solutions. Journal of Inorganic Biochemistry, 2003. 95(1): p. 55–63.

